# Spend today or build for tomorrow? Kinship dynamics and the evolution of alternative helping strategies in cooperative breeders

**DOI:** 10.1101/2025.09.09.675076

**Authors:** DWE Sankey, M Patel, PM Buston, MA Cant, RA Johnstone, T Rueger

## Abstract

Cooperatively breeding animal groups are characterised by the presence of helpers, which assist breeders in raising offspring through a variety of behaviours, such as provisioning young or defence against predators. While these systems have been studied to investigate how costly forms of help can evolve, there is little theory explaining the considerable variation in helping strategies observed across such societies. Here, we use mathematical models to investigate the evolution of alternative helping strategies. Helpers in our models can allocate effort to immediate, short-term benefits or durable, long-term benefits. We find that allocation depends on the kinship dynamics within the group. Specifically, where helpers are more related to future breeders than current breeders, often because they themselves *become* the future breeder, our model typically predicts greater allocation to durable help. Conversely, immediate help is favoured when helpers are less related to future breeders than to current breeders. We use our model to explore how demographic features impact kinship dynamics within the group, identify which factors select for immediate versus durable help, and which amplify conflict between breeders and helpers over helping strategies. The model is a first step towards explaining variation in types of help observed among cooperatively breeding societies.

## Introduction

Cooperatively breeding animal groups have proved particularly valuable systems to study the evolution of cooperation (Cant, 2012; Taborsky et al., 2021). While much attention has focussed on the question of why some non-breeding individuals have evolved to help breeders to rear offspring (Hamilton, 1964; Kokko et al., 2001, 2002), less attention has been paid to explaining the rich variety of helping behaviours we see in nature, such as nest building (Collias & Collias, 1978), territory defence (Langergraber et al., 2017), cleaning (Yllan et al., 2024) and feeding (Koenig et al., 2016), and the fitness consequences of these different behaviours for both the helper and the helped. Current models of cooperative breeding recognise the diversity of helping behaviours but tend to reduce them to a single variable, helping effort (Emlen, 1982; Kokko et al., 2001, 2002). However, this approach assumes that all helping behaviours would be equally preferable to the helper who bears the costs of help, and to the breeder who receives the benefits — assumptions which are unlikely to be warranted outside of clonal communities lacking genetic conflict. Altogether, factors underlying variation in helping strategies observed in nature remain insufficiently understood.

One useful way of differentiating helping behaviours is by considering the durability of their effects on the productivity of a patch. Some actions—such as provisioning food for offspring (Koenig et al., 2016) or leaving a pheromone to demarcate a territory (Jordan et al., 2010) — can increase the short-term productivity of a territory, for example by boosting offspring survival or deterring threats. However, these *immediate* forms of help have transient effects and may offer little benefit to future inhabitants of the same patch (but see: Kokko et al., 2001). In contrast, investments that improve or maintain the territory itself—such as nest construction (Tello-Ramos et al., 2024), long-term territory management (Rueger, Heatwole, et al., 2022; Yllan et al., 2024), or food caching (Koenig, 1980)—can enhance longer-term patch productivity. These *durable* forms of help can benefit future group members across generations (Lehmann, 2007).

Kin selection theory provides a useful framework to understand how selection shapes the allocation of effort to immediate versus durable forms of help. Here, we adopt a kinship dynamics approach (Croft et al., 2021), which goes beyond static measures of relatedness to consider how an individual’s kinship with its social group changes over time in predictable ways. This temporal perspective is crucial, as our distinction between immediate and durable helping behaviours hinges on their differing impacts across time. We might expect more focus on immediate helping behaviours (for example direct offspring provisioning) in populations where group members are closely related and the probability of inheriting a breeding position is low. This is seen in genetically related cooperatively breeding cichlids (*Neolamprologus pulcher*), where helpers directly tend to the breeder’s offspring (Balshine et al., 2001). By contrast, durable help may evolve more readily in low relatedness groups where the probability of inheritance is high, driven by the future direct fitness benefits to the actor. Since subordinates may inherit breeding status in the future, the costs of this form of helping can be offset by the future benefits to the individual that provides it. Indeed, this is seen in low-relatedness groups of the cooperatively breeding clownfish, *Amphiprion percula*, where helpers rarely provide direct care to the breeders’ offspring (Buston, Bogdanowicz, et al., 2007), and instead invest in territory maintenance (Rueger, Heatwole, et al., 2022), a durable form of help which likely benefits both current and future breeders (Barbasch et al., 2020; Buston, 2004a; Chausson et al., 2018; Rueger et al., 2021).

In this study, we model helper strategies that allocate effort between a relatively immediate and a relatively durable form of help (hereafter referred to simply as ‘immediate’ and ‘durable’). We use a kinship dynamics approach to explore how the demographic features of a population (i.e., survival, migration) can shape the evolution of helping allocation. Further, we consider how helping allocation would evolve if it were under breeder control and investigate the conflict between helper and breeder-controlled allocation. Our aim is to provide theoretical underpinnings for how variable demographics of animal systems could explain the variance in helping strategies observed in nature.

## Model and Results

We model an infinite, asexual population divided into discrete ‘patches’, with two individuals per patch: one breeder and one (queuing) helper. In each time step, helpers allocate a proportion, *h*, of their efforts to immediate help, and the remaining proportion (*1-h*) to durable help (Fig. 1A-C; see also parameter table, Table 1). Both types of help boost the fecundity of the breeder in the patch, both in the current and in future time-steps (denoted by *t*). However, the effects of help decay over time, at a rate that differs between the two types. Immediate help provides a larger initial fecundity benefit to the breeder (*d*_*imm*_) than durable help (*d*_*dur*_) where 0 *< d*_*dur*_ *< d*_*imm*_ *<* 1. However, while initially more profitable, immediate help decays more rapidly across time steps at a rate of (1- *d*_*imm*_)^*t*^, whereas durable help decays at a slower rate of (1- *d*_*dur*_)^*t*^, which makes durable help potentially favourable for future breeders on the patch (Fig. 1A). The decay curves for both types of help are such that the total cumulative benefit each type can provide over time is equal (Fig. 1A, B). This ensures that our results reflect only differences in how help is distributed over time, rather than differences in total benefit. We assume that immediate and durable forms of help contribute independently to breeder fecundity but that both exhibit diminishing returns with increasing investment. To achieve this we scale the benefit of immediate help as 1−*e*^−*h*^ and the benefit of durable help as 1−*e*^−(1−*h*)^. These diminishing returns functions inherently favour mixed allocation strategies, maximising total benefit over time when helpers divide effort equally between the two forms of help (Fig. 1B, C). This captures the idea that it may be beneficial for to divide effort between tasks such as feeding (immediate help) and building or maintaining structures (durable help) (Russell et al., 2010; Wong & Balshine, 2011).

**Table 1.**
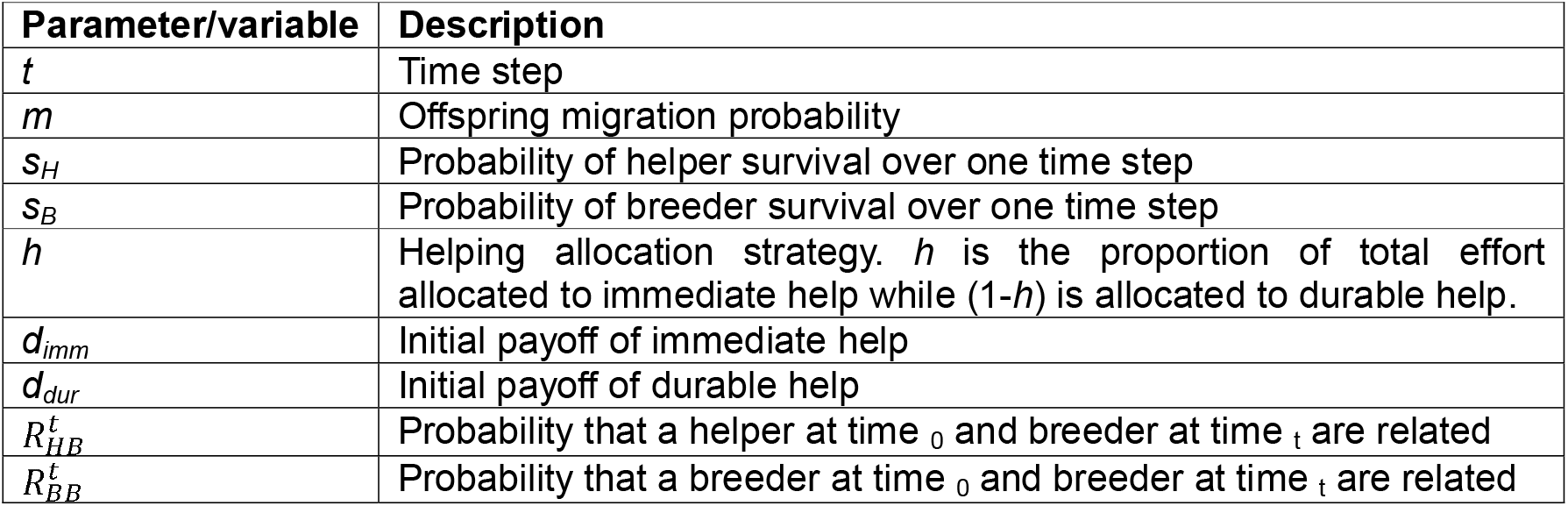
Table of parameters and variables.

| Parameter/variable | Description |
| --- | --- |
| $t$ | Time step |
| $m$ | Offspring migration probability |
| $s_H$ | Probability of helper survival over one time step |
| $s_B$ | Probability of breeder survival over one time step |
| $h$ | Helping allocation strategy. $h$ is the proportion of total effort allocated to immediate help while $(1-h)$ is allocated to durable help. |
| $d_{imm}$ | Initial payoff of immediate help |
| $d_{dur}$ | Initial payoff of durable help |
| $R_{HB}^t$ | Probability that a helper at time $0$ and breeder at time $t$ are related |
| $R_{BB}^t$ | Probability that a breeder at time $0$ and breeder at time $t$ are related |

**Figure 1.**
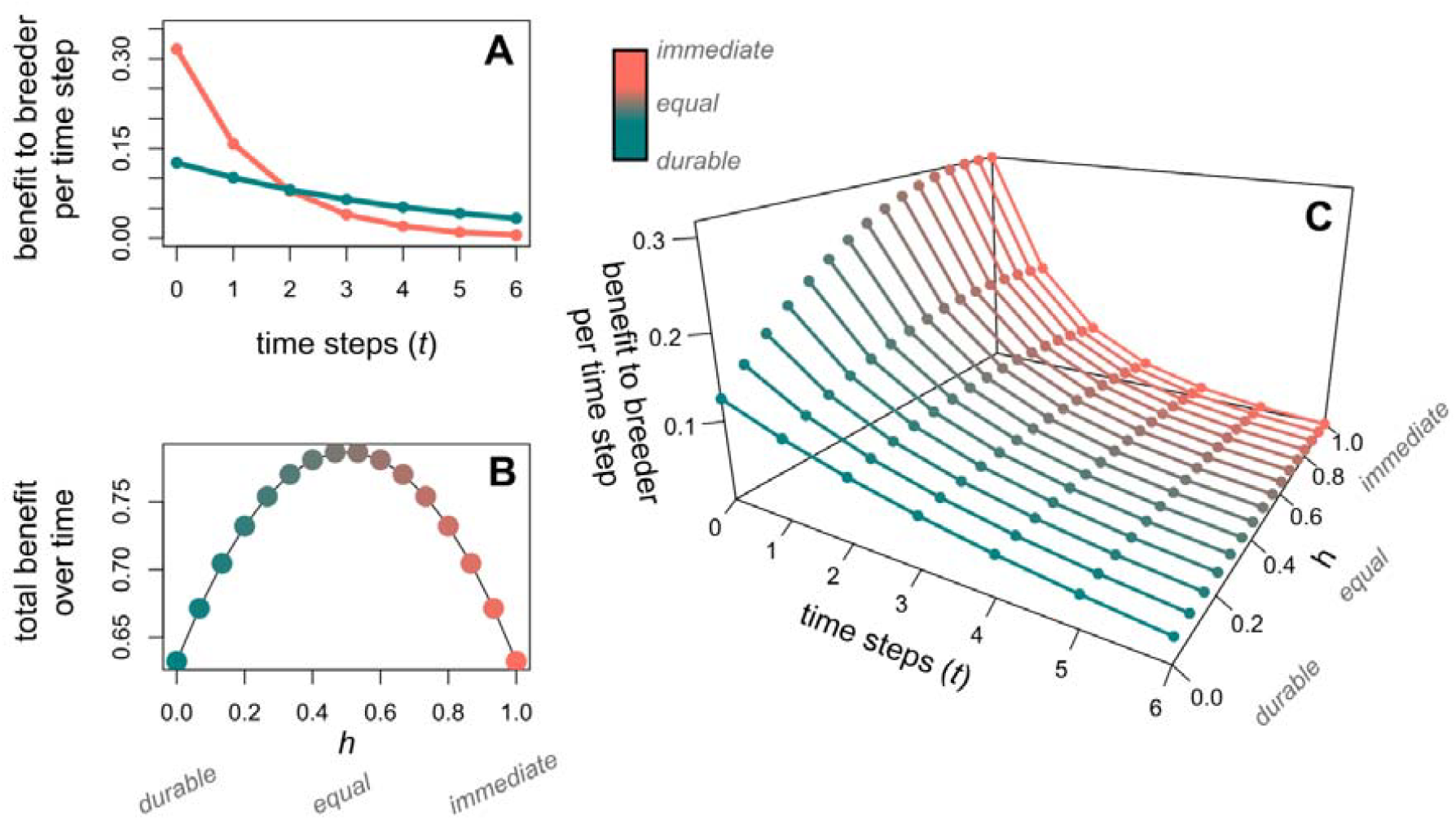
Immediate vs durable helping. **A)**<u>Immediate vs durable trade-off</u>. Fecundity benefit to breeder as a function of time step. In time step zero when the behaviours are performed, immediate help (coral) has a higher potential value to the breeder. However, the value of immediate help decays faster across time steps than the value of durable help (teal). **B)**<u>Summed total benefit</u> to breeders of various helping allocation strategys, *h* (16 representative values). Over time, total benefit to breeders is always higher when helping is allocated more evenly between immediate and durable help. **C)**<u>Helper strategies</u>. Benefit to breeder per time step (*t*), for 16 representative allocation strategies (*h*) of the helper, bounded by the extreme strategies depicted in panel A). Parameters: panel (A) and (C): *d*_*imm*_ = 0.5, *d*_*dur*_ = 0.2; Panel (B) is parameter independent.

In each time step (*t*): the breeder on a patch produces a large number of young, its productivity, *F*_*i*_, dependent on the helping strategies of current and past helpers (Eqn. 1). Then, a fraction, *m*, of the offspring migrate to other patches at random, while a fraction (1 − *m*) remain in the natal patch. A fraction *s*_*B*_ of breeders, and *s*_*H*_ of the helpers, survive to the next time step. If the helper survives and the breeder dies, the helper inherits the patch and becomes the new breeder. Offspring on a patch, both immigrants and locally born, then compete on equal terms to fill any breeder and helper vacancies in the patch (with those that fail to obtain a vacancy of either kind dying). The cycle then repeats.

### Evolution of helping strategies

We use an adaptive dynamics framework to study the evolution of the helping allocation strategy, *h*. In this approach, we assume that the population is almost entirely composed of individuals expressing a resident strategy, with rare mutants differing slightly in their helping allocation strategy. Evolution proceeds by the sequential substitution of rare, small-effect mutations, which sweep to fixation when advantageous. To determine whether a mutant strategy can invade, we assess its fitness when rare in a resident population. To do this, we start by calculating how helping strategies influence breeder fecundity over time, which requires us to calculate the pattern of relatedness between helpers and breeders across time steps (kinship dynamics). We then calculate mutant fitness, by considering how breeders offspring compete locally and globally with other juveniles in the population; which involves determining the reproductive value for both helpers and breeders (see Supplemental Material). Finally, we compute the selection gradient by differentiating mutant fitness with respect to mutant strategy (evaluated at the resident strategy), accounting for the fact that, with probability *r*, its social partners also express the mutant strategy (see Supplemental Material) (Taylor & Frank, 1996). Evolutionarily singular strategies occur where this total derivative equals zero.

### Kinship dynamics

To determine how helping strategies impact the productivity of a mutant breeder (*F□*), we must consider how helping effort was allocated not only in the present time step but across all preceding time steps. This requires calculating the probability that a helper from _*t*_ time steps ago was related to the current breeder. (In our model, individuals are either mutant or resident, so relatedness is either 0 or 1.) If related, the helper would have followed the mutant strategy *h*_*i*_; if unrelated, it would have followed the resident strategy, *h*. Helping effort _*t*_ time steps into the past is given by:

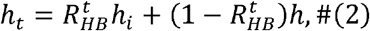

where 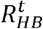 represents “intertemporal relatedness” – the expected relatedness between a helper and a breeder _*t*_ time steps into the *future*. (Importantly, the relatedness of a breeder and the helper on its patch _*t*_ time steps into the past is equivalent to the relatedness of a helper to a breeder _*t*_ time steps into the future. Taking the perspective of helper’s relatedness to future breeders is useful because it is the helper’s strategy which evolves.) In essence, intertemporal relatedness determines the likelihood that the recipient of help will be kin (current and future). It is not explicitly defined in the model but depends on the demographic parameters (*m, s*_*H*_ and *s*_*B*_) of the modelled population (see Supplemental Material for full derivation). For a helper, intertemporal relatedness 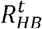 can be lower than that of a helper to the current breeder, 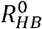. This occurs when population turnover—due to mortality and offspring migration—results in the replacement of the helper’s relatives by unrelated migrants (Fig. 2A). However, intertemporal relatedness can also be greater than their relatedness to a current breeder. This is the case where the helper is likely to survive to *become* the breeder (Fig. 2C).

**Figure 2.**
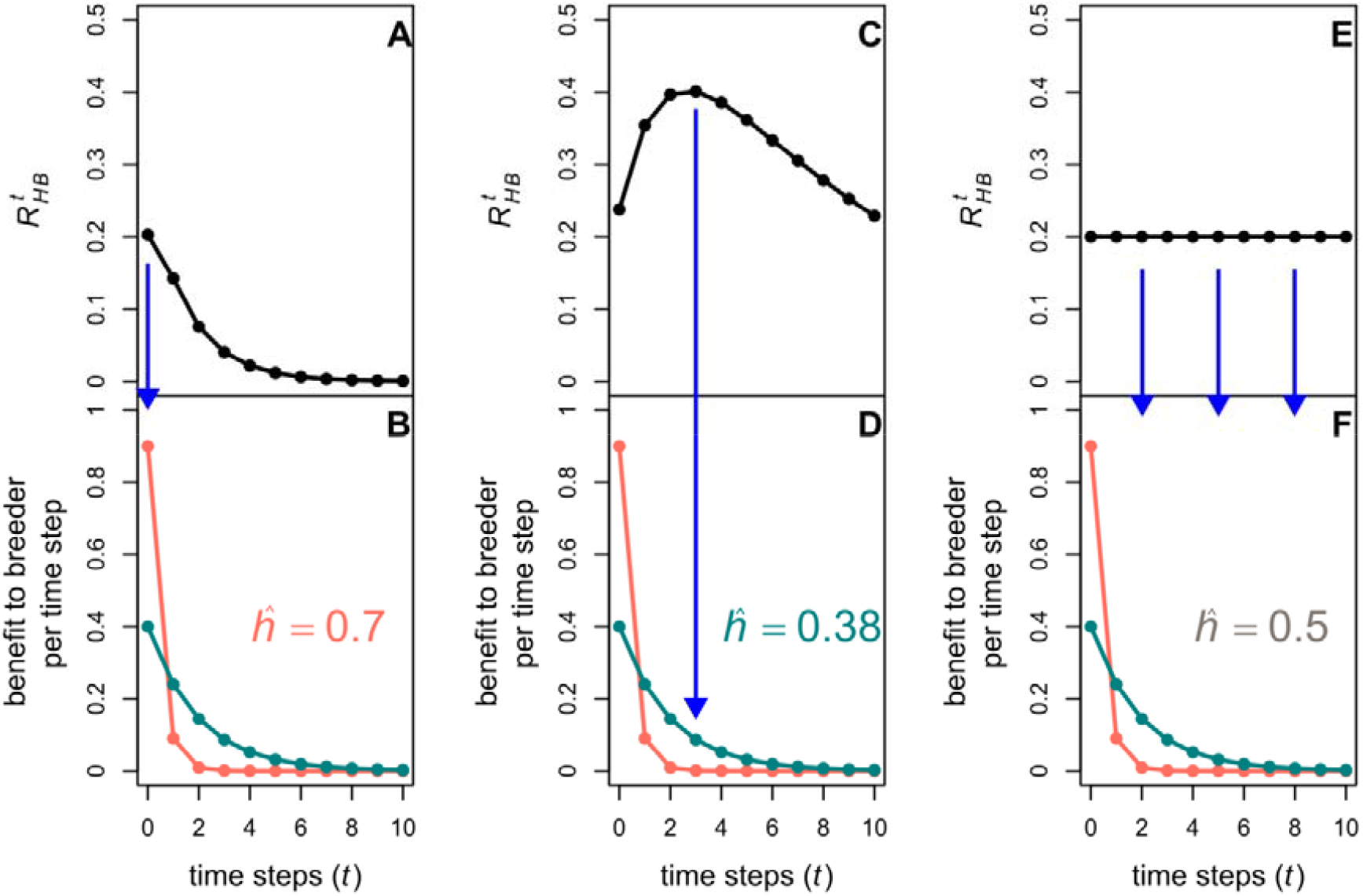
Kinship dynamics and helping allocation. Intertemporal relatedness, (the expected relatedness between a helper and a breeder _*t*_ time steps into the future) is key to determining the evolutionarily stable helping allocation strategy,, because it determines the probability that a relative, or a non-relative will receive the benefit of their help. Panels A, C, E depict over time steps, _t_, for various parameter combinations. Panels B, D and F are similar to Fig. 1A, depicting how fecundity benefit to the breeder decays over time for extreme strategies (*h*=1), coral, representing the decay of immediate benefit, and *h*=0, teal, representing the decay of durable benefit. Parameters for B, D and F: *d*_*imm*_ = 0.9 and *d*_*dur*_ = 0.4. Blue arrows point from the peak of intertemporal relatedness (or multiple time steps in the case with no peak; panels E, F) to the corresponding panels below. They draw attention to the fact that the helping strategy contributing most benefit to breeder at the time step(s) with highest intertemporal relatedness often predicts which strategy will be favoured by selection. **A)** If monotonically decreases, **B)** allocation will always favour immediate benefits (> 0.5). **C)** If, however, relatedness increases (at least for a short period), then **D)** allocation can sway in favour of durable benefits (< 0.5). **E)** Finally, if relatedness neither increases nor decreases over time, intermediate allocation strategies are favoured (, reflecting the benefits of investing in both immediate and durable help. Parameters in panels A-B: *m* = 0.6, *s*_*H*_ = 0.05, *s*_*B*_ = 0.2; in panels C-D: *m* = 0.6, *s*_*H*_ = 0.8, *s*_*B*_ = 0.8; in panels E-F: *m* = 0.8, *s*_*H*_ = 0.8, *s*_*B*_ = 1.

### ESS allocation of help, *ĥ*

Given the fitness of a mutant individual (see Supplemental Material),

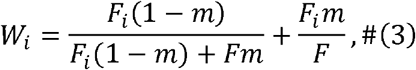

we can solve for ESS levels of *h* (which we denote as *ĥ*) using Taylor and Frank’s (1996) method. We find that the unbounded ESS is

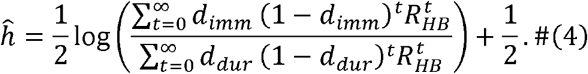

Thus, stable allocation between the two helping behaviours, *ĥ*, depends critically on the ratio between two sums of the geometric series which represent the value of immediate help (numerator), and durable help (denominator), which are multiplied in each time step by intertemporal relatedness for that time step. If the numerator is larger, immediate helping will be favoured in the allocation strategy. Conversely if the denominator is larger, more effort will be allocated to durable help. If the ratio of the sums of the two series is equal to one, then the helping allocation will be equal between immediate and durable help: *ĥ*. This formulation can be viewed as an intertemporal extension of Hamilton’s rule, in which benefits of helping are no longer realised in a single time step, but instead accumulated over time, modulated by how related the future recipients of the help are.

Immediate help is favoured *ĥ* when intertemporal relatedness 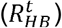 decreases monotonically (Fig. 2A-B). That is, the helper can still allocate some effort to durable help but selection will never favour more durable help than immediate help (Fig. 2B). This is because: while immediate help and durable help are geometric series which both sum to one, immediate helping benefit decays faster than durable benefits, and thus places relatively larger weights on the earlier terms in the series. When 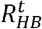 is monotonically decreasing, it is also larger earlier in the series, and so immediate help is always favoured. Note that monotonically declining relatedness is a sufficient, but not necessary, condition for this pattern—non-monotonic declines can also favour immediate help, depending on the relative weighting of benefits across time.

For durable help to be favoured by selection (*ĥ* < 0.5) intertemporal relatedness needs to have a period of increase over time (e.g., Fig. 2C, D). This is possible only when helpers from the present can become the breeders of the future. This is evidenced by the special case where helpers always die (*s*_*H*_ = 0) and so cannot possibly become breeders. Here 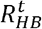 shrinks with each time step: 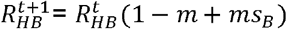.

If intertemporal relatedness remains relatively constant over time, the helping allocation strategy evolves to stability at *ĥ* ≈ 0.5 (roughly equal allocation to both immediate and durable help) (Fig. 2E, F). This reflects the assumed diminishing returns of benefits of engaging in too much immediate or durable forms of investment (Fig. 1B).

### How demographic parameters determine the kinship dynamics

Increasing the probability of helper survival, *s*_*H*_, while keeping all other parameters constant, raises the allocation to durable help because of its impact on the kinship dynamics. This is because, when helper survival is low, monotonically decreases over the time steps, as helpers will be unlikely to become future breeders (Fig. 3A solid line). Accordingly with previous arguments, the stable strategy when *s*_*H*_ is low is to invest more in immediate help. As *s*_*H*_ increases, the helpers have more likelihood of surviving to become the breeder (Fig. 3A, dashed and dotted lines), which raises intertemporal relatedness, and so favours allocation to durable help (Fig. 3A-B).

**Figure 3.**
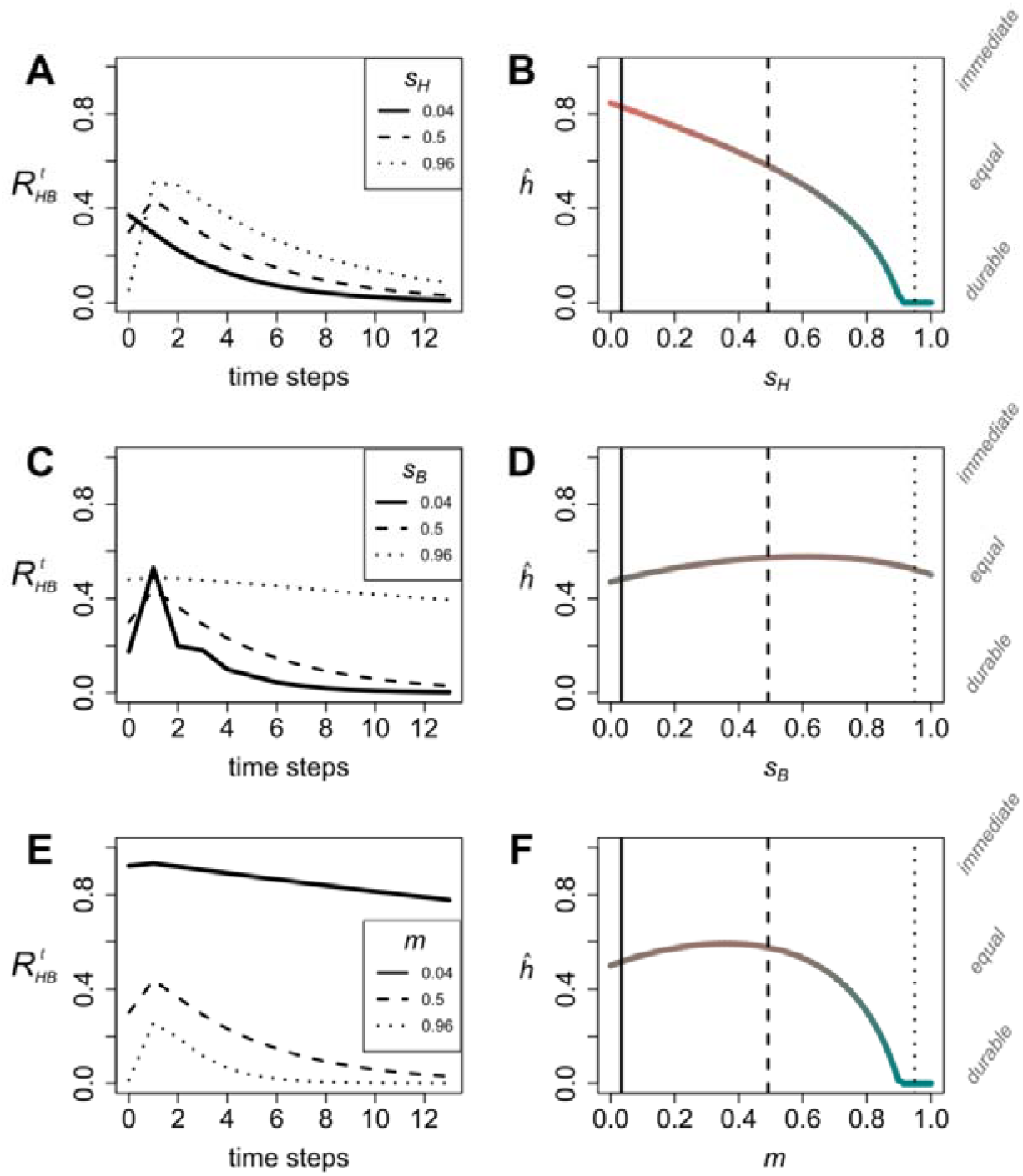
How demography shapes kinship; and how kinship shapes helping. Panels **A–B** show effects of helper survival (*s*_*H*_), **C–D** breeder survival (*s*_*B*_), and **E–F** migration rate (*m*) on (Left) intertemporal relatedness over time and (Right) evolutionarily stable helping allocation. Parameter values given by lines, dots, and dashes on left column (see legend) are matched on the right column. Intertemporal relatedness at time step 0 reflects relatedness between a helper and the current breeder on a patch. Parameters: *d*_*imm*_ = 1, *d*_*dur*_ = 0.2; *s*_*H*_, *s*_*B*_ and *m* = 0.5 unless shown on axes or legend.

When breeder survival rate *s*_*B*_ is high, the breeder is likely to remain in place for many time steps. Intertemporal relatedness remains relatively static (Fig. 3C, dotted line), which favours equal allocation between immediate and durable forms of help (Fig. 3D, also see Fig. 2E, F). Interestingly, equal allocation is also often selected for when *s*_*B*_ is low (Fig. 3C, solid line). Here, although there is an initially large spike in intertemporal relatedness at (favouring durable help), this upward trend is not sustained, because helpers become breeders which are likely to die after only one time step. From this peak onward, intertemporal relatedness generally decreases (favouring immediate help), even though the short-lived breeder’s offspring can cause a second spike a generation later. The balance of these two forces (spike favouring durable; decrease favouring immediate) often leads to an equal allocation strategy (Fig. 3D).

The migration parameter *m* has the largest influence on within-patch relatedness, 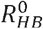. When *m* is low, intertemporal relatedness remains stable and high across time as breeders are likely replaced by their own offspring (Fig. 3E, solid line). Stable intertemporal relatedness favours mid-range allocation to immediate and durable benefits, *ĥ* ≈ 0.5 (Fig. 3F). When *m* is high, within-patch relatedness, 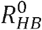, is low. Starting from a low baseline, the impact of helpers surviving and becoming breeders drives an upward trend in intertemporal relatedness (Fig. 3E, dotted line), favouring durable help (Fig. 3F). Yet, for mid-range values of *m* there can be a trend towards more allocation to immediate help. This is because, despite the initial spike in intertemporal relatedness created by helpers surviving to become breeders, relatedness values can still plummet after the spike, creating the decreasing trend in the series over time (Fig. 3E, dashed line) which favours allocation to immediate help (Fig. 3F).

### Mid-range durable benefits favour allocation to durable help

The parameters determining the payoff and decay of immediate and durable help, *d*_*imm*_ and *d*_*dur*_, also impact the evolution of helping allocation strategies. When *d*_*dur*_ is very low, the benefit from durable help is “spread thinly” over many time steps, and high allocation to durable help rarely evolves. Even when intertemporal relatedness initially rises over time – something which usually increases allocation to durable help – relatedness to breeders in the far distant future will eventually decrease (e.g., Fig. 3A, dotted line). Often, this means that the future beneficiaries of this very long-term durable behaviour will be non-relatives, and therefore we find high investment in immediate help when *d*_*dur*_ is extremely low. Conversely, when *d*_*dur*_ is high it resembles *d*_*imm*_, and so we approach equal evolutionarily stable allocation *ĥ* =0.5. It is when *d*_*dur*_ takes intermediate values that we often find the most investment in durable help (Fig. 4).

**Figure 4.**
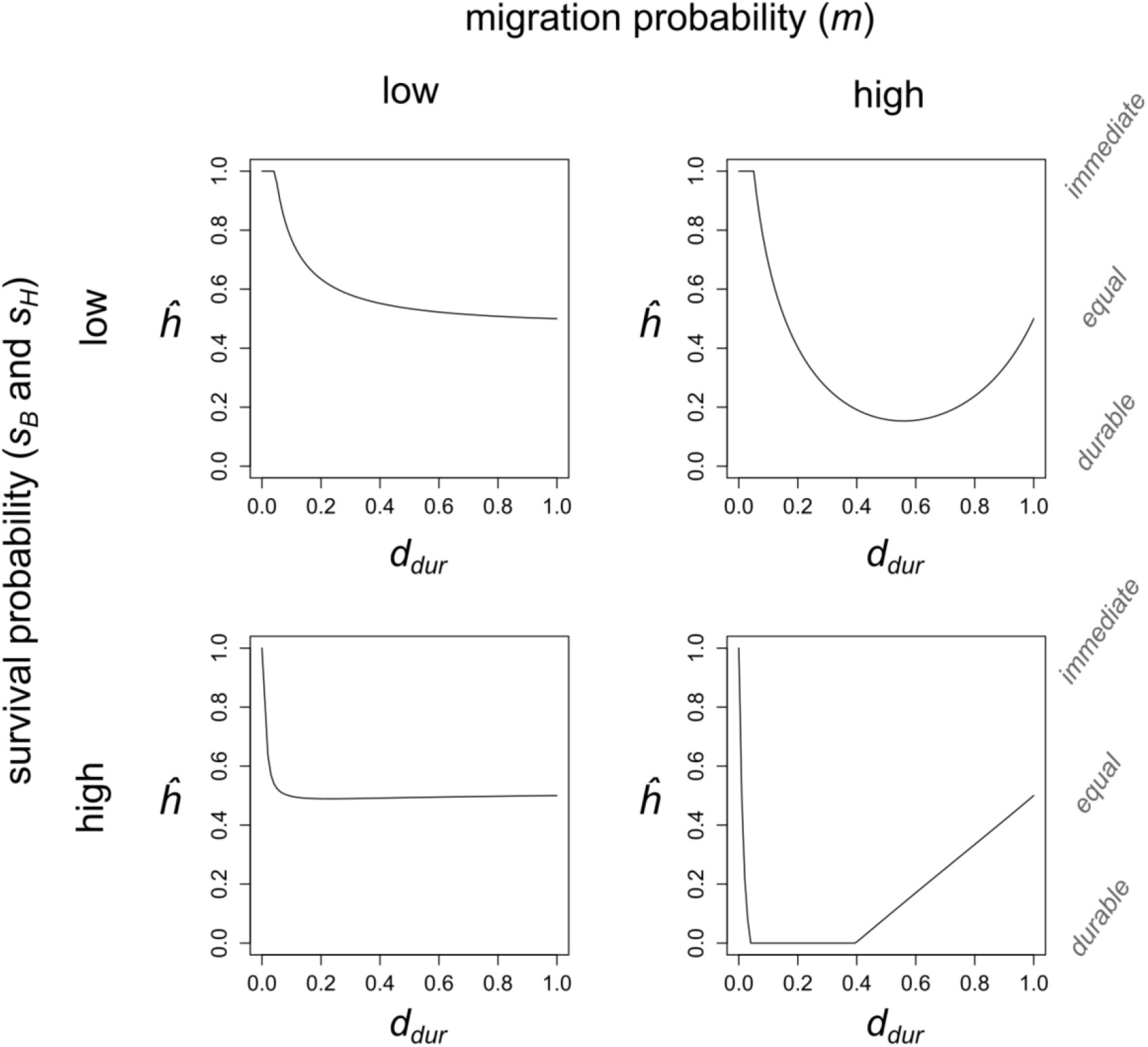
The durability of durable help. Evolutionarily stable helping allocation for migration rates 0.1 (left column) and 0.9 (right column); and survival probability (helper and breeder both equal) at 0.1 (top row) and 0.9 (bottom row). The parameter *d*_*dur*_ determines the initial value of durable help to the breeder in the current time step, but higher initial values come with a cost of faster decay across time steps (lower durability). Across much of the parameter space, mid-range *d*_*dur*_ predicts highest allocation into durable help (low is high allocation to durable help). *d*_*imm*_ = 1 in all panels.

This result – that allocation to durable help peaks at intermediate values of *d*_*dur*_ – depends on the demographic parameters in the model. Firstly, when migration probability is high, we find the relationship more pronounced (Fig. 4). This is mostly because within-patch relatedness is lower, allowing for a more pronounced surge in intertemporal relatedness over time (favouring durable help allocation). Second, when survival probability is high (of both the helper and the breeder) the impact of mid-range *d*_*dur*_ on increasing allocation to durable help is more pronounced and left-shifted (Fig. 4). More pronounced because durable benefits are favoured when helper survival is higher; and left-shifted because if durable benefits are spread across many time steps, higher breeder survival will translate into greater probability that future kin receive the benefits.

### Helping under breeder control

In many systems there is apparent conflict between breeders and helpers over helping behaviour (Reeve & Gamboa, 1987; Reeve & Sherman, 1991). Here we explore potential conflict between breeders and helpers over allocation to immediate versus durable help.

What is the optimal allocation strategy from the perspective of a breeder? That is, what would the breeder choose to do if it could control the allocation strategy,, of the helper? Our model predicts that breeders favour a helper to always invest predominantly in immediate help (excluding extreme parameter values such as *s*_*B*_ = 1, and *m* = 0 where equal allocation to each helping type is preferred). This result arises because the key variable governing their choices is intertemporal relatedness for the breeders 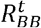. That is, the relatedness between a breeder and a breeder _t_ timesteps into the future. The time series 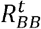 always starts at 1 (breeder is 100% related to itself in the 0^th^ time step), and then decreases monotonically over time, regardless of the demographic parameters (Fig. S1). Since a monotonic decrease in relatedness to future breeders favours immediate help, allocation strategy under breeder control therefore favours more allocation to immediate help than durable help. Helping allocation, *ĥ*, under breeder control also has a simpler relationship with *d*_*imm*_ and *d*_*dur*_: with *ĥ* always increasing for larger *d*_*imm*_ and smaller *d*_*dur*_ (Fig. S1).

### Conflict over helping allocation

Demography has significantly different impacts on helping allocation under breeder control compared to helper control. <u>Migration probability, *m*</u>: higher migration generally increases durable allocation when under helper control, but instead unidirectionally increases allocation to immediate help under breeder control (Fig. 5A). This is because, for breeders, high migration causes a plummet in expected future relatedness (relative to the starting value of 1). When the breeder is unlikely to be replaced by a relative, they favour immediate help. When <u>breeder survival *s*_*B*_</u> is low, breeders favour high investment in immediate help to maximise benefits while they remain the breeder, which conflicts with helpers which invest more in durable because their likelihood of ascension increases. Conversely, when breeder survival rates are high, we see equal allocation of immediate and durable help favoured by both helpers and breeders because the best strategy is to provide a more equal of helping behaviours for the long-lived breeder (Fig. 5B, also see Fig. 2E, F). <u>Helper survival, *s*_*H*_</u> has only a very minor effect on the value that the breeder places on immediate vs durable help (Fig. 5C). This contrasts with the uniformly increasing allocation to durable help for increasing helper survival we saw under helper allocation (Fig. 5C, Fig. 3B). This is because the breeder’s relatedness to future breeders is more influenced by breeder survival (whether it remains breeder) and migration (which implicates the probability it will be replaced by a relative), than whether helpers survive.

**Figure 5.**
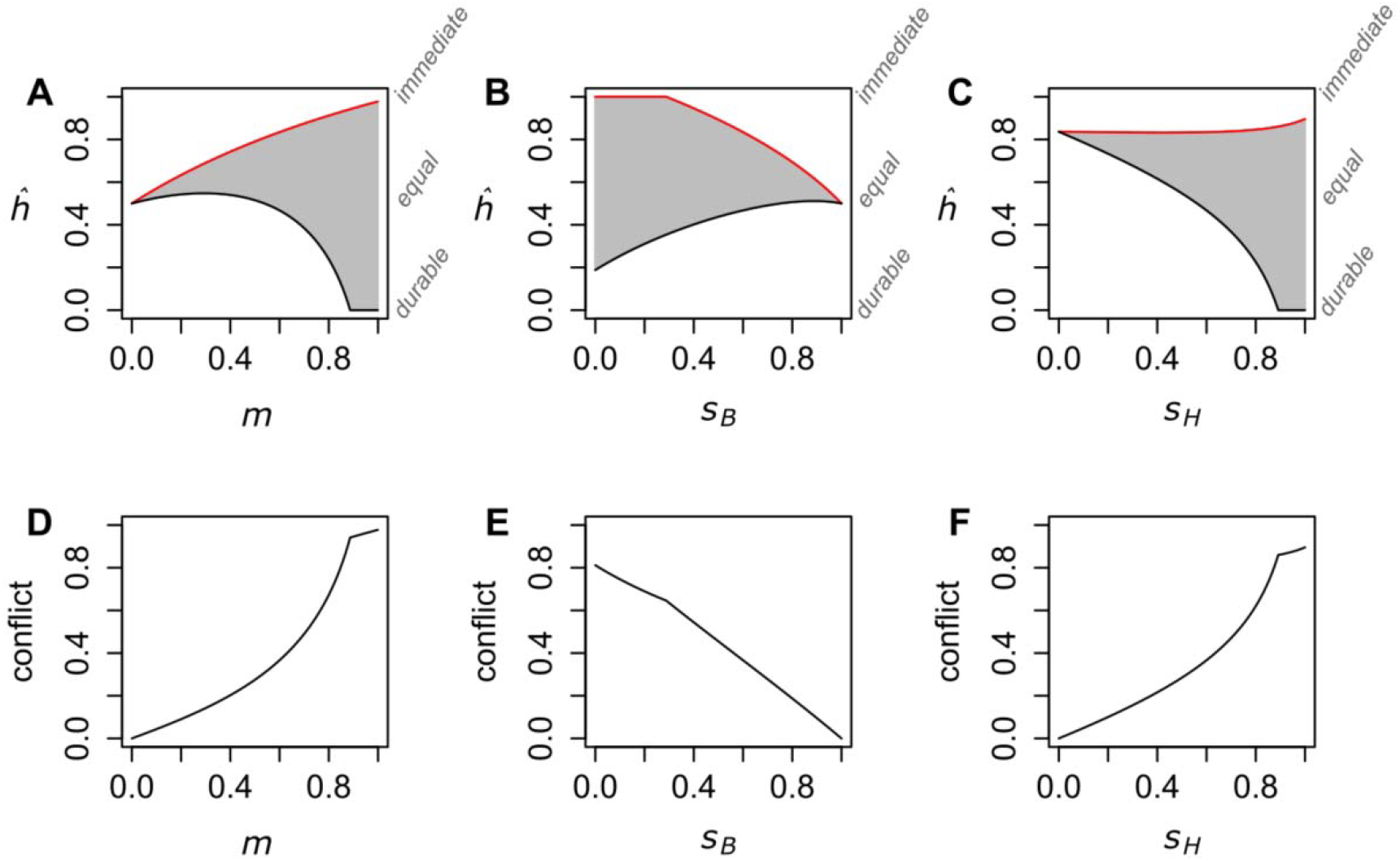
Demographic influences on breeder- and helper-controlled allocation and conflict. Panels A-C). Helping allocation strategy when decision is under breeder control (red), helper control (black) and the zone of conflict (grey) for demographic parameters **A)** offspring migration probability, **B)** breeder survival probability, **C)** helper survival probability. Panels D-F). Conflict (difference between breeder- and helper-controlled allocation) against demographic parameters **D)** migration probability, **E)** breeder survival probability, **F)** helper survival probability. Parameters: *d*_*imm*_ = 1, *d*_*dur*_ = 0.2; *s*_*H*_, *s*_*B*_ and *m* = 0.5 unless shown on axes.

The degree of conflict – which we define here as the absolute difference between helper and breeder-controlled allocation optima (Taborsky et al., 2021) – increases uniformly with increasing migration probability, and helper survival, and decreases uniformly with increasing breeder survival (Fig. 5A-C). These monotonic patterns in conflict contrast with the often non-monotonic effects of demography on allocation strategies under helper and breeder control (Fig. 5A-C). For conflict, monotonic trends held across the full parameter space, confirmed by differentiating conflict with respect to each parameter and evaluating all parameter combinations from the interval [0.01, 0.99] sampled at five evenly spaced points (e.g., see: Fig. 5D–F).

## Discussion

Our results demonstrate that helping allocation can be shaped by the relatedness between helpers and the future occupants of the breeding position on their patch. When helpers are more likely to be related to future breeders than to current ones—particularly when they have a high chance of surviving to inherit the breeding position themselves—they often favour investment in durable help. Conversely, when helpers are more weakly related to future breeders than current ones, immediate help is favoured. Thus, the kinship dynamics over time—not simply relatedness at a single point—emerges as a key driver of variation in helping strategies. This is contrary to the superficially plausible expectation that greater instantaneous (within-patch) relatedness between the helper and breeder should select for more allocation to immediate forms of help. High within-patch relatedness *can* favour more allocation to immediate help, since high relatedness to the current breeder implies that the relatedness to future breeders is unlikely to be higher – though this is not always the case. For example, see Fig. 2A, C, for a case of two systems where the population with higher within-patch relatedness has greater allocation to *durable* help. These results provide insights into well-studied cooperatively breeding animal systems and provide predictions for future research into breeder/ helper conflict.

While caution should be applied in overinterpreting the model’s relevance to empirical patterns before formal tests are conducted, patterns across cooperatively breeding species broadly align with the model’s predictions linking kinship dynamics to helping strategies. In meerkats (*Suricata suricatta*), helpers are highly related to current breeders but less so to the future breeders in their natal group due to frequent dispersal (Clutton-Brock et al., 2001; Clutton-Brock & Manser, 2016; Maag et al., 2018), and they predominantly perform forms of immediate help such as brood care and sentinel duty (Clutton-Brock et al., 2004). Meerkat burrows might be durable structures, but meerkats tend to shift breeding location often (Strandburg-Peshkin et al., 2020), and so there is perhaps limited “storing” of breeding value within the territory for future generations. Naked mole-rats (*Heterocephalus glaber*), with low migration and stable relatedness to future breeders due to rare breeder turnover, exhibit a seemingly more equal mix of both durable (burrow maintenance) and immediate (food sharing, defence) helping behaviours (Sherman et al., 1992). Sociable weavers (*Philetairus socius*) also show mixed helping effort: while breeder turnover is more frequent than in mole-rats, both high local relatedness and common ascension, factors which lead to equal allocation in the model, are present in this species. Accordingly, sociable weavers’ exhibit both high investment in both durable help (communal nest construction) and immediate help (offspring provisioning) (Covas et al., 2006; Covas & Du Plessis, 2005; van Dijk et al., 2014). Finally, clown anemonefish (*Amphiprion percula*) show little immediate help but invest in durable territory maintenance, consistent with high expected relatedness to future breeders due to queuing and territory inheritance (Barbasch & Buston, 2018; Buston, 2004b, 2004a; Buston, Bogdanowicz, et al., 2007; Rueger, Heatwole, et al., 2022).

More explicit efforts to measure the intertemporal relatedness (the expected relatedness between helpers and future breeders) across a whole range of taxa will be essential to test the generality of our model. For example, recent work in seven group-living mammal species shows that local relatedness can change systematically with age and that these patterns can be predicted from basic demographic parameters (Ellis et al., 2022). Subsequent research should also isolate relatively durable and relatively immediate helping forms and estimate time budget allocation to these tasks.

Our analysis focuses solely on the fitness incentives of helpers and breeders in an idealised population, omitting many physiological, morphological, and (much of the) ecological constraints that are likely to limit the forms of help across species. For example, cooperatively breeding white-fronted bee eaters (*Merops bullockoides*) nest in sandbank cavities prone to rapid environmental degradation (Wrege & Emlen, 1991), which may influence their investment in these structures. In other cases, morphological or physiological limitations may constrain what forms of help are feasible. Phylogenetic analyses could be useful for disentangling the relative influence of these factors (Gonzalez-Voyer & von Hardenberg, 2014). Nevertheless, these drivers are not mutually exclusive from inclusive fitness maximising incentives, and our model provides a foundation for understanding variation in helping behaviours across populations.

### Conflict

Our model raises questions over breeder helper-conflict. While it may seem reasonable to assume that helping decisions are more under helper control than breeder control, this may not always be the case. For example, breeders might act to modify the behaviour of the helpers. For example, with the threat of eviction which can induce helpers to “pay to stay” on the patch (Zöttl et al., 2013, but see: Cant, 2021). Similar arguments have been made regarding who has control over helper reproduction (e.g., see: Buston, Reeve, et al., 2007; Buston & Zink, 2009; Fischer et al., 2014; Johnstone, 2000; Johnstone & Cant, 1999; Reeve et al., 1998; Reeve & Ratnieks, 1993). Much of the existing discussion has focused on breeders forcing helpers to increase effort, but it may be just as important to consider whether they are coerced to change how that effort is allocated. The key question may not be whether helpers work, but where they invest their efforts.

We found two clear predictions for the degree of conflict between breeders and helpers over helping allocation. Firstly, populations with higher probability of offspring migration would have greater conflict over helping allocation. Second, populations with higher probability of helper survival relative to breeder survival have greater conflict over helping allocation. (In all cases of conflict, breeders are predicted to prefer greater allocation to immediate benefits.) But how can we test these predictions regarding conflict over helping allocation? One simple and useful measurement could be the rate of agonistic interactions between breeders and helpers which might be expected to increase with increasing the demographic parameters which predict conflict. Also revealing would be how breeders allocate their own time when helpers are experimentally removed could potentially reveal the types of help they value in the absence of the helper. Perhaps most compelling might be to observe how breeders respond to helpers with different levels of helping allocation, and to find some way of experimentally manipulating the types of help that helpers provide. Some tests may be more species specific, but also more revealing of the underlying conflict (Taborsky et al., 2021).

### Further implications

Our results urge us to consider an active role of mutualistic partners. We found was that when durable investments return value at a steady rate (neither too short-term nor too stretched across multiple time steps), helpers are often incentivised to invest more in these resources with steady returns. This raises a provoking possibility for species in which durable help entails cooperative interactions with interspecific mutualists (Buston, 2004a; Chausson et al., 2018; Kautz et al., 2009; Rueger, Heatwole, et al., 2022). If we assume the mutualist fitness increases with increasing helper allocation into durable help, it could pay the mutualist to control the rate of return on helpers’ durable investments to a “not too fast; not too slow” rate which maximises future investment in the mutualist. This idea aligns with empirical examples where mutualists shape the reward structure to filter and condition partner behaviour. For instance, in acacia–ant mutualisms, plants secrete sucrose-free nectar that only their defensive ant partners can metabolize (Kautz et al., 2009). This effectively ties the mutualist ants to the host and helps prevent exploitation by non-defenders, illustrating how mutualists can manipulate the payoff environment to maintain cooperative investment. In an anemonefish-anemone mutualism, fish vociferously consume nutritious egesta released from the anemone (Rueger, Bhardwaj, et al., 2022; Verde et al., 2015), a resource whose release rate could plausibly be regulated by the anemone itself. Future models could incorporate active mutualists and coevolving strategies to better understand the ecology of durable helping behaviours in mutualisms— covering a broad range of obligate and facultative mutualisms.

## Conclusion

Our model provides a new theoretical framework for understanding variation in helping behaviours in cooperatively breeding systems. By distinguishing between immediate and durable help and linking their evolution to kinship dynamics across time, we show that relatedness to future, not just current, breeders is key to predicting helping strategies. This approach generates testable predictions about demographic and ecological conditions favouring different types of help, and highlights overlooked sources of conflict between breeders and helpers. It offers a foundation for future empirical and theoretical work on the diversity of cooperative strategies observed in nature.

## Supplemental Material

### Kinship dynamics derivation

When a mutant helper allocates *h*_*i*_ to immediate help and (1-*h*_*i*_) to durable help, then allocation at time _0_ is given by,

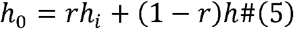

where *r* is relatedness between the current breeder and helper. To calculate *F*_*i*_, we need to understand how helping efforts were allocated not just at time _0_ but in *all* time steps prior to _0_. For this, we need to calculate the probability that the helper any n time steps ago was of the mutant stain. We first calculate relatedness at demographic equilibrium at time _t_, and then work backwards.

First, let *r* denote the expected relatedness between the breeder and helper on the patch. Then the equivalent value in the next time step, *r’*, is given by,

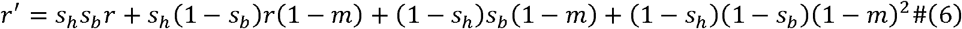

To see where this comes from, with probability *s*_*H*_*s*_*B*_, both the helper and the breeder survive, and relatedness between the pair, *r*, remains unchanged; with probability *s*_*H*_(1-*s*_*B*_) the breeder dies and the former helper survives to become the new breeder: in this case, with probability (1 − *m*) the helper vacancy is filled by a locally-born offspring of the former breeder, related to the new breeder on average by *r*; with probability (1 – *s*_*H*_)*s*_*B*_ the former helper dies, and, with probability (1-*m*), the new helper is an offspring of the breeder, and therefore related by a coefficient of 1 (because reproduction is asexual); finally, with probability (1-*s*_*H*_)(1-*s*_*B*_), both the breeder and helper die. Then with probability (1 − *m*)^2^ both are locally born offspring of the former breeder. By setting *r’* equal to *r* we then solve for the expected relatedness at demographic equilibrium.

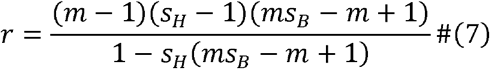

Recall that the amount of benefit provided to the breeder at time _t_ is dependent its relatedness to past helpers, because relatedness to past helpers determines the historical allocation of *h*_*i*_ vs *h* on their patch. For this relatedness over time – or, kinship dynamics – we define a new nomenclature: firstly, 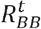 representing the relatedness between a focal breeder and a breeder, n time steps into the future. The current breeder might be related to the future breeder either because they are siblings, the future breeder is the current breeder’s offspring, or the future breeder could even *be* the current breeder: if it survives across the _t_ time steps. Similarly, 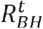 represents the relatedness between a focal breeder and future helper; 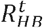, the relatedness between focal helper and future breeder which occupies the future patch; and 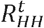, the relatedness between a focal and future helper. At time _t_ the relatedness vector *v*^*t*^ is given by 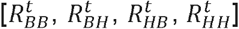. To calculate *v*^*t+1*^, we multiply *v*^*t*^ by the transition matrix, *A*.

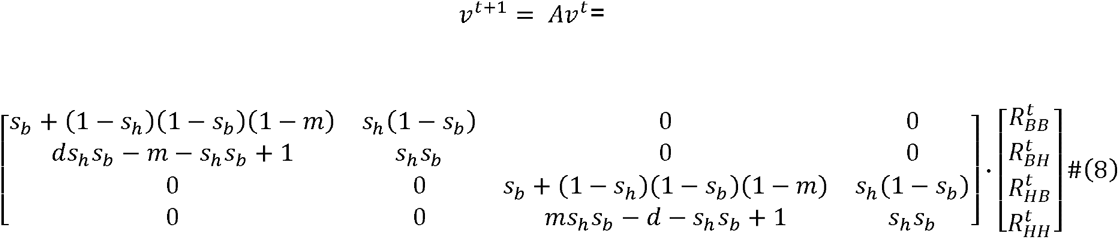

To illustrate this process, we detail how to update 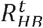 to 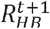. Other updates follow a similar process. 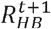 is the probability that the helper at time _t_ and the breeder at time _t+1_ are related. There are two avenues by which 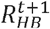 can be non-zero. Firstly, there is a chance, *s*_*H*_(1- *s*_*B*_) that the helper at time _t_ and breeder at time _t+1_ are related because the time _t_ helper *is* the breeder at time _t+1_. This is because the breeder dies (1- *s*_*B*_) and the helper survives (*s*_*H*_), and transitions to breeder. To update this first possibility *s*_*H*_(1- *s*_*B*_) is multiplied by 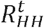. (See third row; fourth column of *A*). Secondly, there is a chance [*s*_*B*_ + (1- *s*_*H*_)(1- *s*_*B*_)(1-*m*)] that 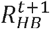 is nonzero because the helper and breeder were already related at time _t_. This is because, with probability *s*_*B*_, the breeder survives and 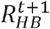 is equal to 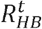. It does not matter whether the helper survives, *s*_*H*_, or dies (1- *s*_*H*_), because we are calculating the relatedness between the time _t_ helper and the time _t+1_ breeder; finally, with probability (1- *s*_*H*_)(1- *s*_*B*_) both the helper and breeder die. In this case, 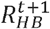 is equal to 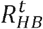 only if the new breeder is the breeder’s offspring, which happens with probability (1-*m*). To update this second possibility 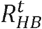 is multiplied by (*s*_*B*_ + (1- *s*_*H*_)(1- *s*_*B*_)(1-*m*)) (See third row; third column of *A*).

Given that *F*_*i*_ depends on help accrued from 0 to ∞ time steps ago. We need to then calculate *v*^*t*^ for any _*t*_ time steps. We do this by first substituting *A*^*t*^ with its diagonalised form:

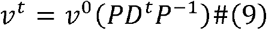

Because we know *v*^*0*^ is equal to [1, *r, r*, 1], we then can calculate (for any *s*_*H*_, *s*_*B*_, and *m*) precisely the probability that current breeder is related to the helper which lived _t_ time steps ago, 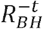, which is equal to 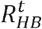, (the third element of the *v*^*t*^ vector). Knowing the probability that the helper _t_ time steps ago was related also means knowing the probability the allocation of help was provided by an individual of the mutant strain.

### Fitness derivation

The fitness of a mutant is given by

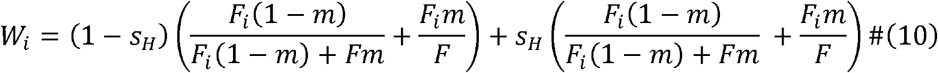

This is because fitness depends not just on fecundity, but also *i*) the probability that those offspring fill a breeder or helper position, *ii*) the breeding values of breeders and helpers (breeding value represents the average number of rounds of breeding an individual achieves) and *iii*) the fair ‘lottery’ for breeder and helper positions among resident and immigrant offspring individuals. Firstly, breeder spots are available with probability (1- *s*_*H*_)(1- *s*_*B*_), and the breeding value of a breeder is 1 + *s*_*B*_ + *s*_*B*_^2^ + *s*_B_ ^3^ … which simplifies to 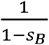. One round of breeding is a given from their first time step as a breeder, and then following this they have *s*_*B*_ chance of surviving each into each subsequent time step for another round of breeding. Multiplying the breeding value of breeder by availability of breeder positions gives a breeding value of (1- *s*_*H*_). If competing locally, the *F*_*i*_(1−*m*) who do not migrate compete fairly with the *Fm* immigrant individuals. Those *F*_*i*_*m* who do migrate compete with the population average number of offspring, *F*. Secondly, the term on the right-hand side describes the availability and breeding value of helper positions. Positions are available with probability 1- *s*_*H*_ + (1- *s*_*B*_)*s*_*H*_: that is, the probability the helper dies, and breeder dies and helper inherits breeder position respectively. The breeding value of a helper is the probability they transition to a breeder, (1- *s*_*B*_)*s*_*H*_ + *s*_*H*_*s*_*B*_(1- *s*_*B*_)*s*_*H*_ + (*s*_*H*_*s*_*B*_)^2^(1- *s*_*B*_)*s*_*H*_ … which simplifies to 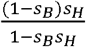, multiplied by the breeding value they attain after they become a breeder 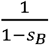. Altogether, this simplifies to *s*_*H*_, the probability of helper survival. Again, a fair lottery decides whether immigrant or local individuals attain the vacant position. Helper survival *s*_*H*_ can be removed from the fitness equation to give the form presented in the main text (Eqn. 3).

### Additional plots

**Figure S1.**
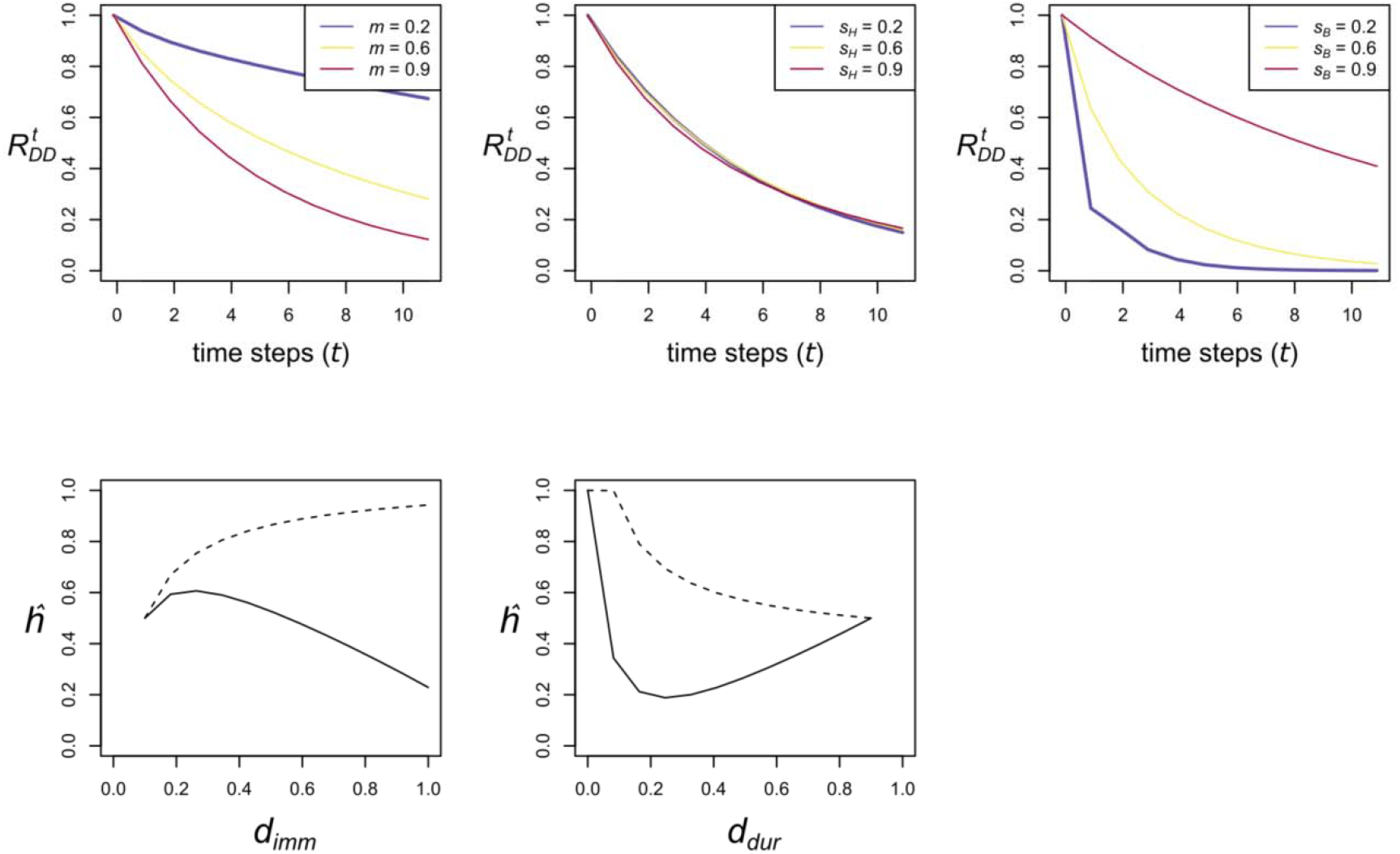
(Top row). The relatedness of a breeder to the breeder, _*t*_ time steps into the future. (Bottom row). Helping allocation, *h*, as a function of *d*_*imm*_, where *d*_*dur*_ is set at 0.1 (left) and *d*_*dur*_ where *d*_*imm*_ is set at 0.9 (right). Solid line = helper. Dashed line = breeder. Parameter values *m, s*_*H*_, and *s*_*B*_ are all equal to 0.8 unless given in legend.

## Notes

### Competing Interest Statement

The authors have declared no competing interest.

